# Unique PKD like protein kinase in *Entamoeba histolytica* regulates vital actin mediated processes

**DOI:** 10.1101/677500

**Authors:** Mithu Baidya, Santoshi Nayak, Amit K. Das, Sudip Kumar Ghosh

## Abstract

*Entamoeba histolytica* causes widespread amoebiasis in humans. Multiple lines of emerging evidence have identified a repertoire of proteins involved in the process of erythrophagocytosis. However, the early initiation of the erythrophagosome at the site of erythrocyte’s attachment is not well understood. Here in our study we have identified and characterized a small Protein kinase D like protein (EhPKDL) in *Eh*, which nucleates actin polymerization and thus mediates many vital processes in *Eh* including erythrophagocytosis. Following multiple biochemical and biophysical approaches, we have characterized EhPKDL and have shown that EhPKDL can indeed interact and prime the nucleation of monomeric actin for polymerization. Furthermore, we went on to demonstrate the vitality of the EhPKDL in major actin-mediated processes like capping, motility, and erythrophagocytosis following knockdown of EhPKDL in the cellular context. Our study thus provides novel insights into the early actin nucleation in *Eh* and thus bridges the gap with our previous understanding of the assembly of erythrophagosomes.

## Introduction

Protein Kinase D (PKD) is implicated in diverse cellular processes that include vesicle trafficking, cell motility, cell proliferation, and apoptosis by participating in distinctly different cell signaling pathways (1, 2). PKDs are Ser/Thr protein kinases, and the subfamily contains three members *viz*. PKD1, PKD2, and PKD3. Though PKD catalytic domain has some similarities with PKCs, it has substrate specificity quite different to that of PKCs and is not inhibited by PKC specific inhibitors like GF109203X and Go6983, Ro-31-8220, nor is it downregulated by phorbol esters, as in case of PKCs (3). In fact, PKDs relay signals downstream after being activated by Protein kinase Cs (PKC). PKCs activate PKDs by phosphorylation of serine residues in the activation loop of the kinase domain (4, 5).

Alternatively binding of diacylglycerol (DAG) in the cysteine-rich domains of PKDs also can stimulate PKD activation (6). Also, Gβγ (G protein) complex can activate PKD1 by interacting with the Pleckstrin homology (PH) domain directly. Thus, the mode of activation of PKDs restricts its response to specific organelles such as Golgi complex, nucleus, plasma membrane, or the cytoplasm (6). The versatility of PKD lies in the multiplicity of its substrates, which it chooses, in particular, circumstances. In the Golgi complex, PKD acts on the oxysterol-binding protein (OSBP) (7), phosphatidylinositol 4-Kinase III-β (PI4KIIIβ) and ceramide trafficking protein (CERT) (8) proteins, which regulates fission of vesicles that are flagged for the plasma membrane. PKD regulates various transcription factors like runt-related transcription factor 2 (RUNX2) and myocyte enhancer factor-2 (MEF2) by phosphorylation of class IIa histone deacetylases (HDAC) (9, 10). Emerging details are delineating PKDs’ role in the regulation of actin dynamics at multiple levels. PKDs presides over actin dynamics by directly modulating regulators like E-Cadherin, cortactin, slingshot protein phosphatase 1 (SSH1L), SNAIL (a transcription factor), Ras and Rab interactor 1 (RIN1), p21-activating kinase 4 (PAK4) and few others (11). In fact, G protein-coupled receptor activation can induce a rapid translocation of PKD1 to the plasma membrane (12). But PKD activity needed for F-actin stabilization, essential for the extension of dendritic spines was demonstrated by Bencsik et al., (13). Though the above study exhibits the involvement PKD in the regulation of actin dynamics, the direct interaction of PKD with actin has never been reported. Therefore, the present study, we report a small primitive PKD like kinase (PKDL) from *E. histolytica* that nucleates the G-actin monomer for polymerization and thus drives all the vital cellular processes of this amoeba involving actin dynamics.

## Results

### *Entamoeba histolytica* contains a small length Protein kinase D like kinase

Analysis of *E. histolytica* kinome reveals the presence of many unusual kinases, very distinct and distantly related to their counterparts from higher eukaryotes. Many of the parasitic kinases possess high sequence similarity in the region of the catalytic domain of known protein kinase subfamily, but they lack many non-catalytic domains characteristic of such kinases in higher eukaryotes. Such kinases are abundant in *E. histolytica*. Interestingly, few kinases of PKA/PKG-like subfamily member has Pleckstrin homology (PH) domain, but there is no PKD identified. In the present study, we have identified a small PH domain-containing protein kinase with the catalytic domain being very similar to PKD kinase (Fig. 1A) as evident from the phylogenic analysis (Fig. 1B) of the protein sequence with known PKDs from many kinases. PH domain similarity amongst all the known PKD members ranges from 34-61%, and the catalytic domain seems to be more conserved (60-93%) (5) and EhPKDL shows significant similarity well within this range, with all the known PKD members but is found most close to PKD1 subgroup of lower vertebrates (Figure S1). Therefore, the protein was tested for phosphotransfer activity in histone type III protein, a universal protein kinase substrate. The recombinant EhPKDL cloned in pGEX-4T1 and expressed in *E. coli* BL21 (DE3) and purified (Fig. 1C), has successfully phosphorylated the histone type III protein suggesting the protein as an active protein kinase. There was no significant inhibition by both genistein (Tyrosine kinase inhibitor) and staurosporine (Ser/Thr kinase inhibitor) like other known PKDs (Fig. 1D). The enzyme kinetics of EhPKDL estimates the *V_max_*of the enzyme as 93.24 nmol/min/mg and *K_m_* value as 4.78 ± 0.32 µM for ATP (Fig.1E). Localization of EhPKDL with specific custom made antibody shows the protein to cytosolic like any other PKDs (Fig.1F). Thus above results suggest that the PH domain-containing protein kinase, so far unannotated, with proven kinase activity and deeper sequence analysis shown to be similar to that of human and mouse Protein kinase D (PKD) (Fig. 1A and B). Therefore, now onwards it will be referred to as Protein kinase D like kinase (PKDL) in the text.

**Figure. 1.**
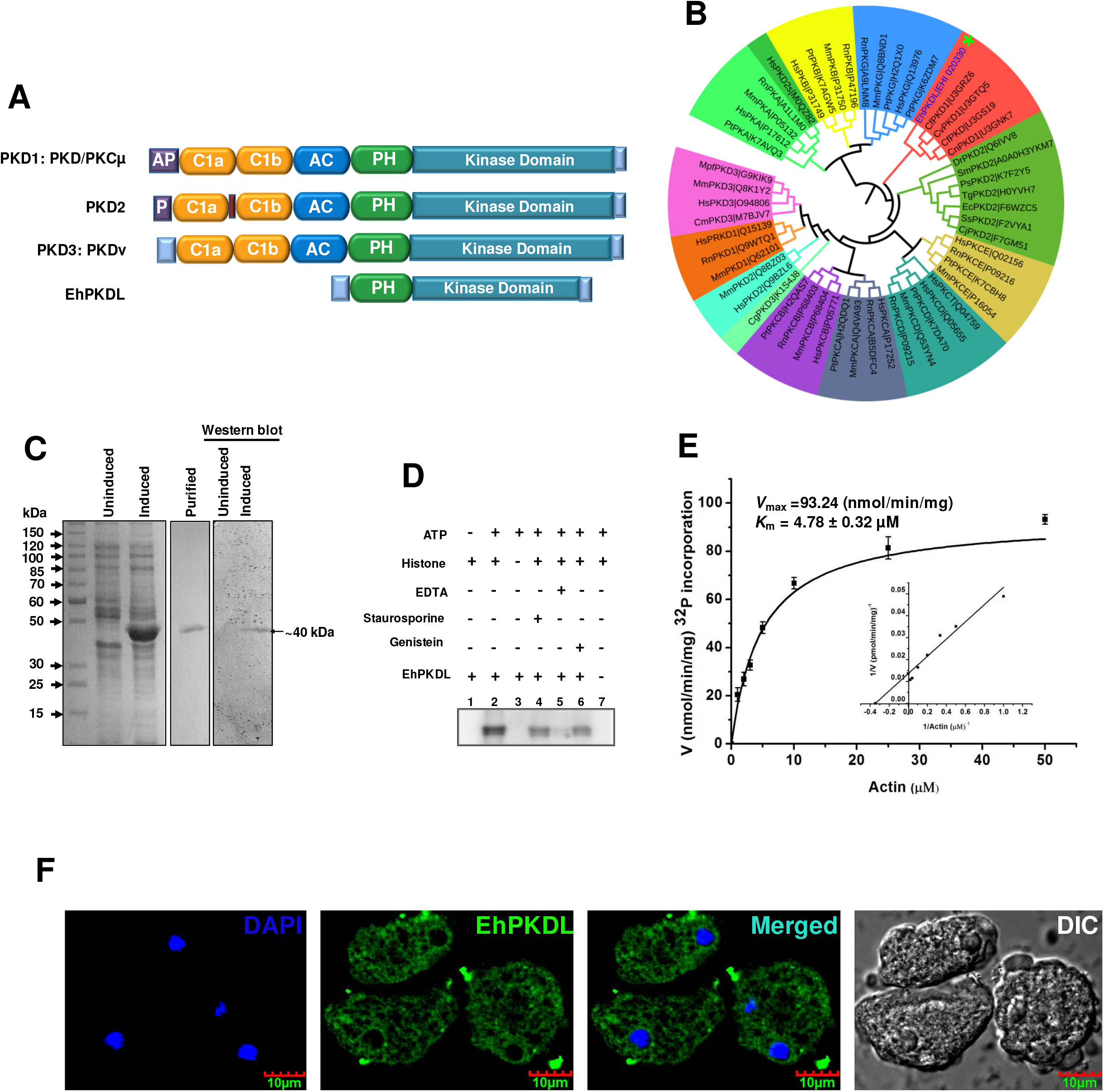
Characterization of a novel PKD like kinase in *Entamoeba histolytica*. A. Domain architecture of various Protein kinase D vis-à-vis known and classified PKDs. B. Phylogenetic tree of EhPKDL developed after comparison with various protein kinases including PKA, PKB, PKC (including all sub groups) and PKG from various higher eukaryotes. (Abbreviation used for the organisms are listed in a separate section).C. Expression of EhPKDL in BL21(DE3) *E. coli* strain cloned in pGEX-4T-1vector. Induced lane contains expressed EhPKDL protein stimulated with 0.2 mM IPTG for 16 h at 16°C, as compared to uninduced lane; lane M contains protein molecular weight marker. Purified EhPKDL protein shown is obtained after glutathione affinity purification and gel filtration. Western blot analysis of uninduced and induced EhPKDL expressing bacterial cell lysate by EhPKDL specific antibody, raised in rabbit.D. γ-^32P^ phosphotransfer activity of EhPKDL with calf thymus histone type III-A as substrate carried out at 30°C shows successful phosphotransferase activity even in the presence of Staurosporine (Ser/Thr protein kinase inhibitor) and Genistein (Tyr protein kinase inhibitor). E. Enzyme kinetic study of EhPKDL. Michaelis– Menten and Lineweaver–Burk plots were used to quantify its kinetic parameters, *V_max_*, and *K_m_*. F. Localization of EhPKDL in *Eh* cells with a specific polyclonal antibody developed in rabbit shows the presence of EhPKDL mostly in the cell cytoplasm.

### Actin is the substrate of EhPKDL

Protein phosphorylation is an important mode of coded messaging employed for cell signaling. Phosphorylation by upstream kinases to a specific downstream protein will actually determine the type and magnitude of the response. Therefore, identification of downstream substrates is central to decipher the molecular function of the protein. To identify the substrate of the EhPKDL, specific antibody raised against EhPKDL was used to pull down the native EhPKDL from cell lysate by Protein A beads. The idea was to pull down all the EhPKDL interacting proteins from *Eh* cell lysate. All the interacting proteins were resolved in SDS-PAGE, and a comprehensive mass spectroscopic analysis was carried out to identify all the proteins after tryptic digestion. Matrix-assisted laser desorption/ionization-time of flight (MALDI-TOF) mass spectroscopy, enabled the identification of actin as one of the interacting protein (Figure S2A and B).

The affinity of the interaction between EhPKDL and actin were further confirmed by pull-down assay, where His-tag actin was immobilized on to Ni-NTA beads as bait, and subsequently, GST-EhPKDL with varying concentration was incubated. Finally, His-tagged proteins, along with the interacting partners, were then eluted by imidazole. On SDS-PAGE analysis, EhPKDL was found in the eluted fraction along with His-tagged actin. Moreover, the interaction was found to increase with increasing EhPKDL concentrations (Fig. 2A). To further substantiate EhPKDL interaction with actin, we resorted to fluorescence spectroscopy, which can also throw some light on the stoichiometry of interactions. Here, EhPKDL-FITC was the donor molecule that on titration with actin-TRITC showed a gradual increase in fluorescence quenching. Since enzyme-substrate interaction is considered a dynamic process, hence the fluorescence quenching observed was also considered as collisional quenching. Collisional quenching of fluorescence is described by the Stern-Volmer equation, where quenching data are usually presented as plots of F0/F versus [Q]. On fitting the quenching data in the Stern-Volmer equation, it generated a straight line, which indicates a single kind of tagged molecule in which fluorophores are equally accessible to the quencher. From Hill’s plot, the stoichiometry of interaction between EhPKDL-actin was determined as 1:1 in a molar ratio (Figure S3A-D). To ascertain if the interacting actin is the substrate for EhPKDL, phosphorylation reaction was carried out as previously described for histone type III. After kinase assay with actin as substrates, a successful phosphotransfer activity was shown (Fig. 2B), with *V_max_* of 120 nmol/min/mg and *K_m_* is 2.93±0.15 µM (Fig. 2C).

**Figure. 2.**
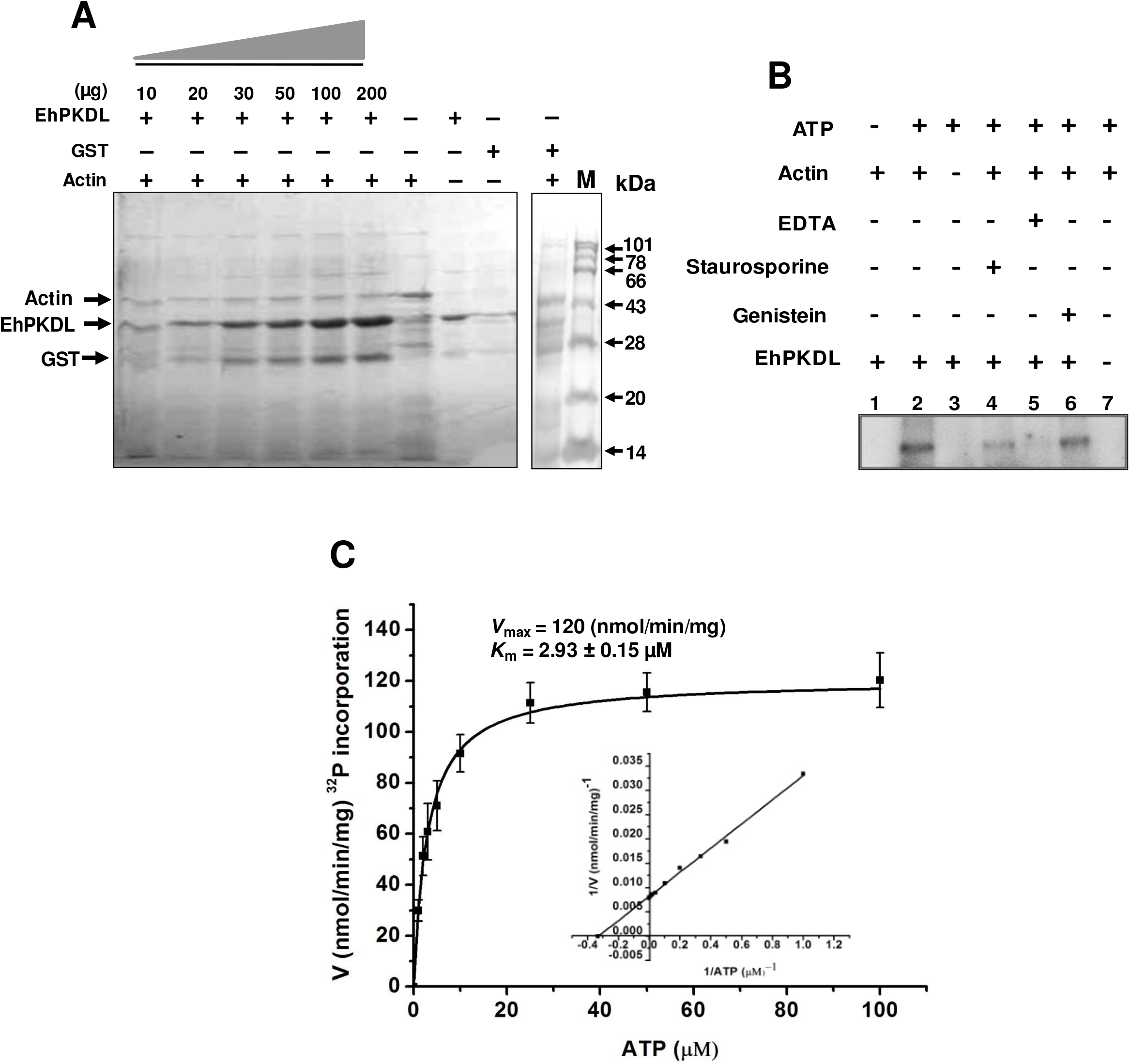
Actin is the substrate of EhPKDL. A. Pull-down assay of EhPKDL with actin. 6xHis-actin (10 µg) was immobilized as bait, to which varying concentration of GST-EhPKDL was allowed to interact. Finally interacting protein complex were eluted and analyzed by SDS-PAGE, which affirms interaction between Actin and EhPKDL. B. Autoradiograph showing γ-^32P^ phosphotransfer activity of EhPKDL with actin as substrate carried out at 30°C, which confirmed phosphorylation of actin by EhPKDL. C. Finally, in enzyme kinetic study of EhPKDL Michaelis–Menten and Lineweaver–Burk plots were used to quantify *V_max_* and *K_m_*of EhPKDL with the substrate as actin.

### EhPKDL primes the nucleation of actin polymerization

Since actin was identified as one of the substrates, which by its polymerization dynamics manifests its function as a seasoned cytoskeletal protein of the cell, therefore, we investigated if EhPKDL has any role to play in actin dynamics. Actin has an inherent property to self-polymerize that seems to accelerate in the presence of EhPKDL but not GST, which suggests EhPKDL helps in polymerization (Fig. 3A). Since a G-actin binding motif was identified in EhPKDL (^89^NVLFSADFEETSALGVYQSTVE^111^) (Fig. 3D), therefore, we analyzed its binding pattern with G-actin by fluorescence spectroscopy. The monomeric pyrene G-actin usually has low signal. Even when EhPKDL was introduced in the reaction with pyrene G-actin at a 1∶1 molar ratio of actin to G-actin binding motif, there was no visible spectrum change as actin remained in its G configuration. But with the subsequent stepwise increase in G-actin titrant, an appearance of the first F-actin signal was observed which we presume may be due to saturation of all the actin-binding domains, which we believe becomes due to dimerization of EhPKDL, as evident during the gel filtration elution profile (Fig. 3C). With the further addition of unlabeled actin, F-actin signal begins to increase when this primed pyrenylated G-actin in actin binding motif (Fig. 3D) help transform G-actin into its F-actin conformation with stepwise integration of G-actin into its nucleus (Fig. 3B). There by, the above data suggest that actin is initially primed in actin binding domain that then gets phosphorylated and subsequently absorbed into the evolving chain of F-actin filament that emerges from it.

**Figure. 3.**
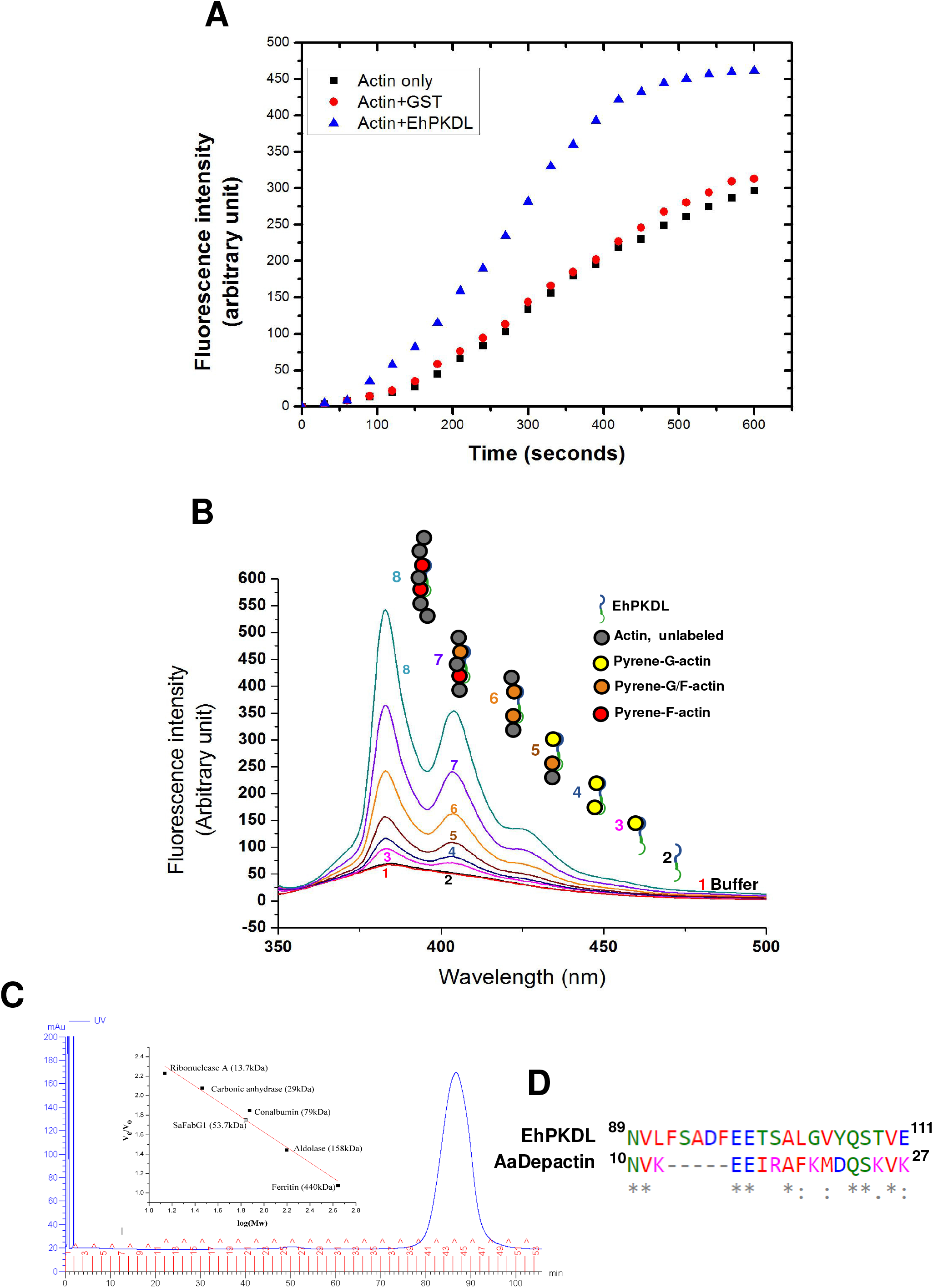
EhPKDL nucleates actin. A. 3 µM of actin was polymerized, either alone (red) or in the presence of 2.5 µM EhPKDL (blue) or with only GST (black), where the highest amount of polymerization was seen in the presence of EhPKDL. B. In actin nucleation study initially, four molar excess of pyrene actin was added stepwise, followed by the addition of unlabeled G-actin. Red spectra correspond to reaction only having polymerization buffer. Addition of 0.2 μM EhPKDL (black), did not show any spectral change until 0.2 μM pyrene actin (pink, blue), which increased fluorescence in expected line due to the presence of pyrenylated G-actin. But subsequent titration with 0.4 μM aliquots of unlabeled G-actin elevated the pyrenylated F-actin signal without the further introduction of any pyrene actin (brown, orange, purple, cyan), indicates integration of G-actin in the growing F-actin filament. The cartoons at the right depict the proposed mechanism of the stepwise evolution of an F-actin nucleus along with the actin-binding domain of EhPKDL. C. Gel filtration profile of EhPKDL in S200 sepharose column. Elution peaks at the 42nd fraction, which from the calibration graph indicates spontaneous dimerization of EhPKDL. D. G-actin binding site of EhPKDL, which is found significantly similar to *Asterias amurensis* (starfish) Depactin’s actin-binding protein.

### EhPKDL modulates erythrophagocytosis

The efficient phagocytic activity of *Eh* cells determines its virulence. *In vitro* phagocytic activity of *Eh* cells is often measured by its ability to engulf human red blood cells (RBC). Upon RBC binding to the cell membrane, phagocytic cup grips around the RBC and then it pulls in the cup into the cytoplasm forming a mature phagosome. These structural changes are mediated by the formation of F-actin wreath around the phagosome, which supports the structure (14, 15). To assess the influence of EhPKDL in F-actin formation around the phagocytosing RBC, which subsequently determine its rate of erythrophagocytosis; *Eh* cells, treated and untreated with EhPKDL specific dsRNA (Fig. 4A and 4B) were fed with human RBCs. The cells treated with EhPKDL dsRNA showed three times less erythrophagocytosis compared to untreated cells (Fig. 4C and 4D). Moreover, EhPKDL treated dsRNA cells scored 3 phagocytic cups at 5 min and 7 cups at 10 min per 50 cells which is significantly less than the control and BS1 (non-specific) dsRNA (treated cells (Fig. 4E). To study the distribution of EhPKDL during erythrophagocytosis, cells were stained for F-actin with phalloidin (red), and EhPKDL was stained with the specific antibody (green). Confocal images clearly indicate that EhPKDL, which is cytosolic in nature, relocalizes around the newly formed cup decorated with actin, soon after the attachment of RBC to the *Eh* (Fig. 4F & I). Interestingly, however, as the cups mature into a phagosome, EhPKDL withdraws from the mature phagosome and is not observed to further co-localize with F-actin (Fig. 4F, panel 2 and 3 and 4H). The above observation goes on to suggest that EhPKDL is required for phagosome formation where F-actin assembly is needed to support the newly formed cup.

**Figure. 4.**
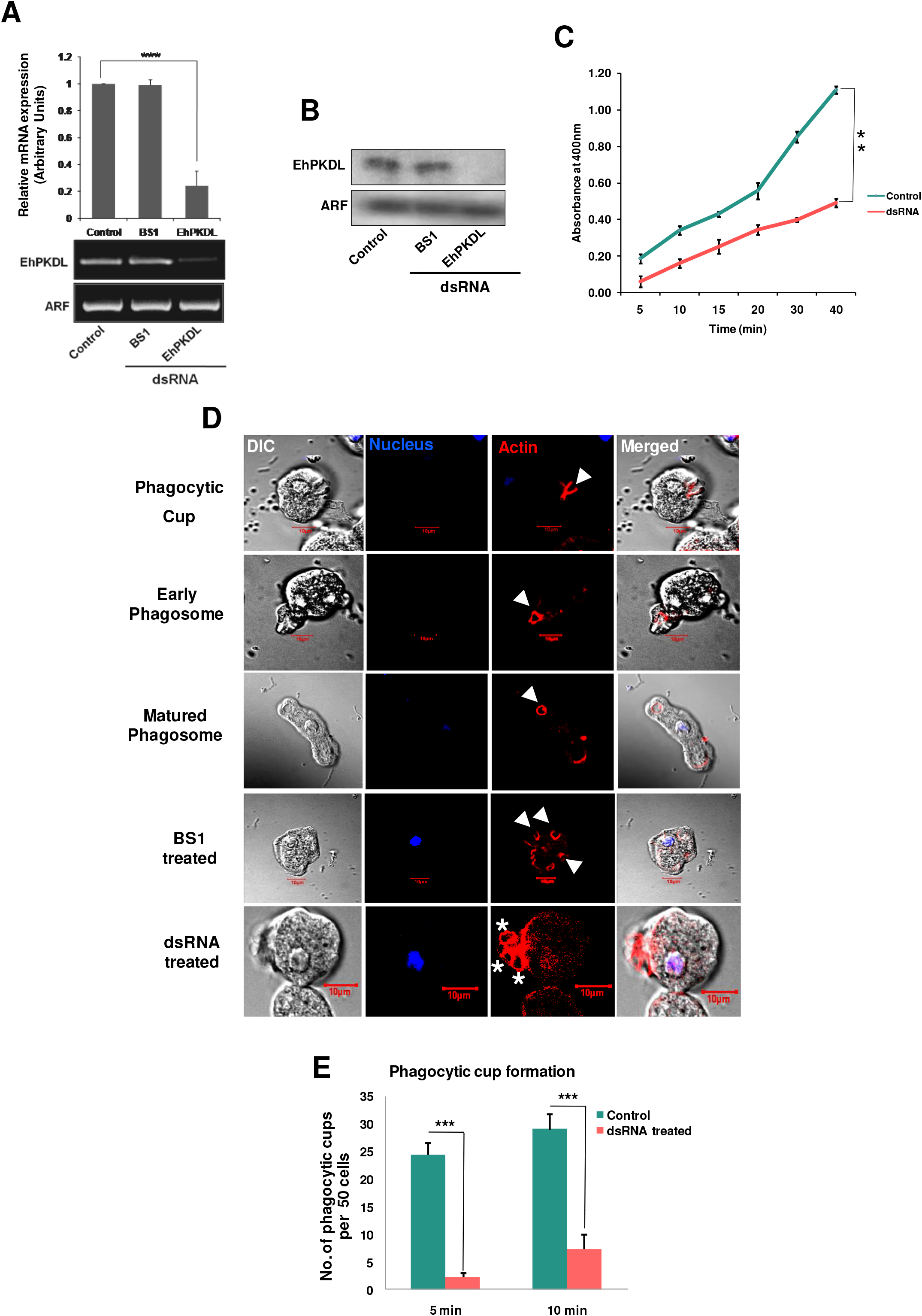
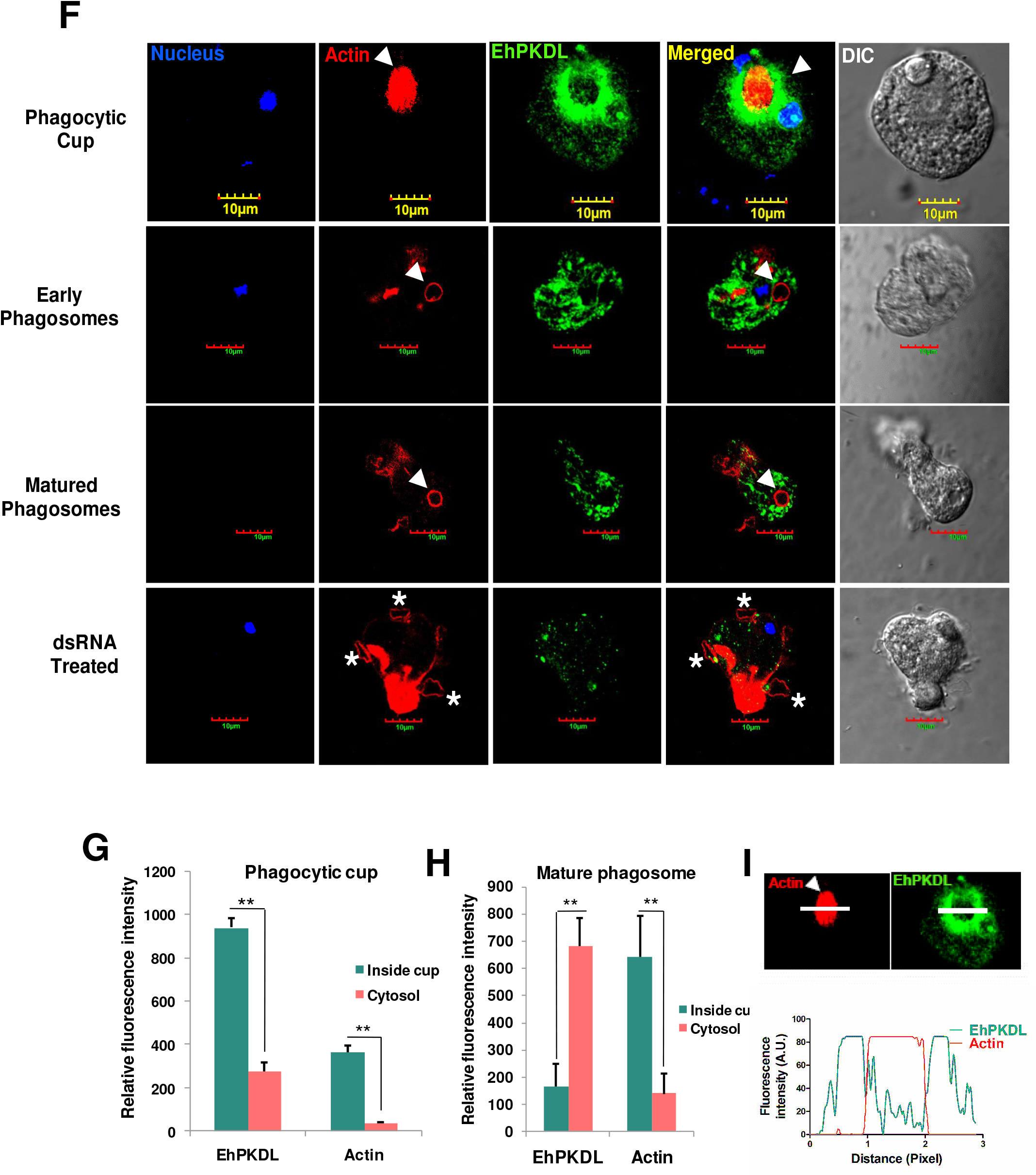
EhPKDL regulates erythrophagocytosis. Down-regulation of EhPKDL. A. sqRT-PCR analysis and B Western blot reveals a significant decrease in relative EhPKDL mRNA expression on EhPKDL specific dsRNA treatment compared to control and non-specific dsRNA (BS1). ADP-ribosylation factor (ARF) gene was considered as the internal control. C. RBC phagocytosis assay identifies a drastic reduction in erythrophagocytosis by *Eh* cells soaked with EhPKDL specific dsRNA. The experiments were repeated three times in triplicates. One-tailed Student t-test was used for statistical comparisons (**p-value≤0.01). D. *Eh* cells treated with or without dsRNAs for 24 h were allowed to engulf RBCs for different timings at 37°C so to capture different stages of phagosome formation. Cells were then fixed and stained for actin with TRITC-Phalloidin, and nucleus with DAPI. Polymerized F-actin at phagocytic cups is marked by solid arrowheads. Asterisks show attached but not engulfed RBCs. Cells treated with EhPKDL specific dsRNA was unable to form proper phagocytotic cups required for engulfment of RBCs. E. For estimation of phagocytic cup formation quantitatively, fifty cells were randomly selected for each experiment as indicated, the number of phagocytic cups present at 5 and 10 min was accessed. One-tailed Student t-test was used for statistical comparisons. ***p-value≤0.005. Statistical data indicates a significant reduction in cup formation at 5 mins, and by 10 mins slight increase in the number of cups infers that for some cells if not complete inhibition but the delay was ensured in the formation of the cup. F. For localization of EhPKDL in RBC engulfing cells, fixed cells were immunostained with anti-EhPKDL antibody followed by Alexa-488, DAPI staining for the nucleus, and actin was stained with TRITC-Phalloidin. Micrograph indicated colocalization of EhPKDL with polymerized F actin at early cup formation but not at later stages. Quantitative estimation of localization, extrapolated from fluorescence intensity, of F-actin and EhPKDL in phagocytic cup and cytosol in G initial phagocytic cup and H mature phagosome. I Line scan analysis by ImageJ across the phagosome.

### EhPKDL regulates cap formation

Capping is an immune evasion mechanism employed by *E. histolytica*, wherein any external molecule that comes and binds to *E. histolytica* surface proteins gets pooled and aggregated around the uroid region to subsequently fall off, thereby inhibiting further immune augmentation. This selective congression of surface-bound protein with the external agents is mediated by actin (16). EhPKDL is expected to influence capping, but to gauge its primacy we have induced capping in *Eh* upon ConA binding to surface glycoproteins. Capping reduced from 70% to 12% in EhPKDL specific dsRNA treated cells in comparison to untreated cells (Fig. 5A and 5B). Confocal micrograph images show that uncapped cells have surfaced bound ConA all over but could not successfully form a cap. On localization study, actin and ConA showed significant co-localization in the cap but with very little presence of EhPKDL (Fig. 5C), thus indicating EhPKDL withdrawal from F-actin once it has evolved from G-actin monomer.

**Figure. 5.**
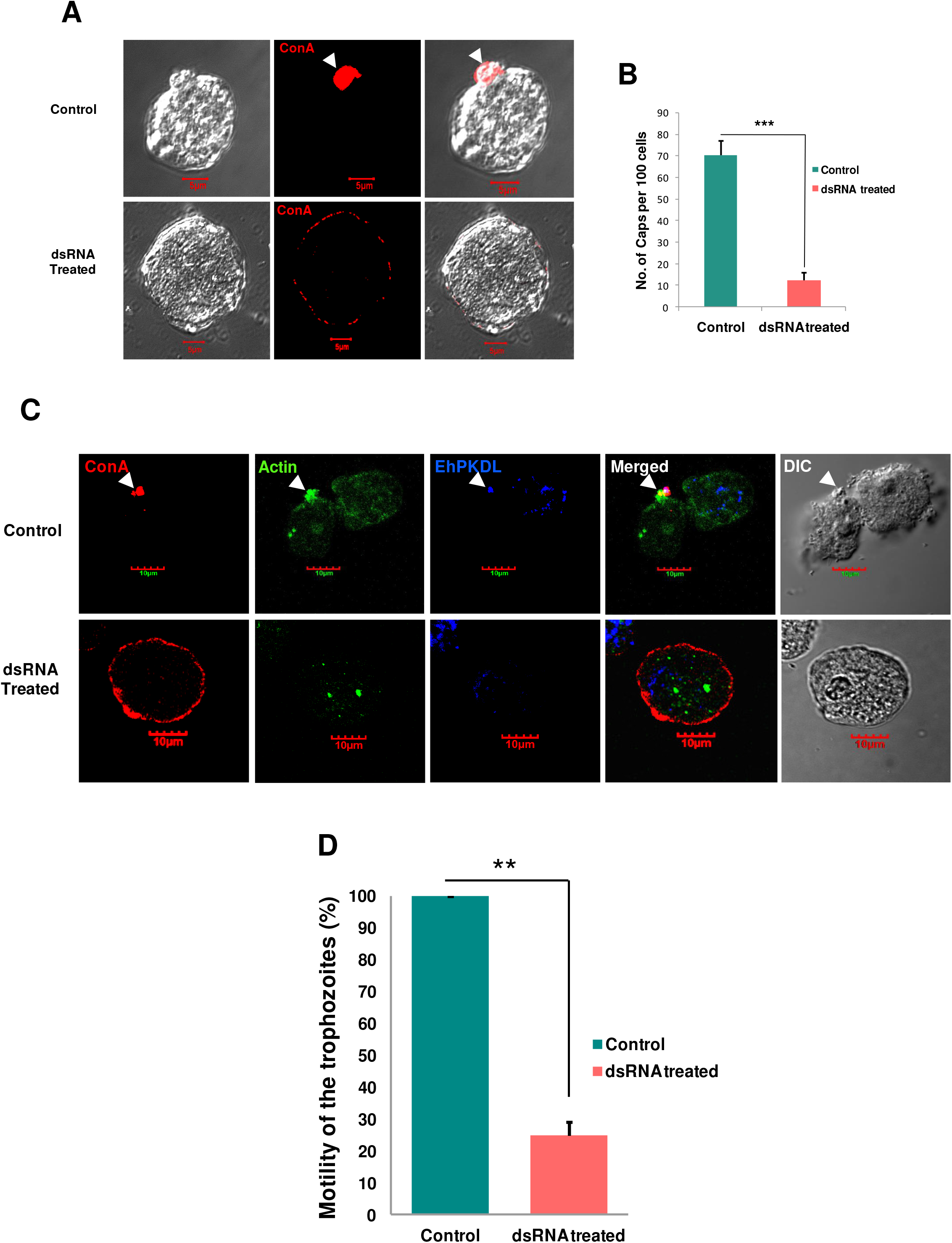
EhPKDL regulates cap formation and motility. A. For induction of cap, TRITC-ConA was allowed to bind the cell surface glycoproteins (as indicated), and cap formation was induced. In untreated cells, caps were observed, but in EhPKDL dsRNA treated cells, ConA remained uniformly bound to the cell surface without any aggregation. Caps are marked by a solid arrowhead. B. Quantitative estimation of cap formation from randomly selected 100 cells indicated a drastic reduction in cap formation in EhPKDL specific dsRNA treated cells. C. For co-localization studies, cap induced cells were immunostained with anti-EhPKDL antibody followed by Alexa 405 tagged secondary antibody, actin was stained with Alexa 488-Phalloidin. On microscopic study, it was observed that EhPKDL co-localized in the cap, but in the EhPKDL dsRNA treated cells it did not; also, the treated cells were observed with reduced F-actin formation. D. The motility of *E. histolytica* trophozoites in control and dsRNA treated cells were investigated using the transwell assay (for details check materials and methods). The motility of control trophozoites was considered 100%. The motility of the dsRNA-treated trophozoites was significantly reduced from that of the control. Mean ± standard deviation are calculated from three independent experiments. An unpaired Student’s t-test was performed for statistical comparison. **p-value≤0.01.

### EhPKDL is stress responsive

Since PKDs are known to respond to stress (17), we checked if EhPKDL has any role to play in stress signaling in *Eh*. *E. histolytica* cells were exposed to different kinds of stress, and their relative mRNA expression was gauged by RT-PCR. *Eh* cells seem to over express EhPKDL in all kinds of stress, but their maximum expression was found to be in heat stress, followed by oxidative stress induced by H_2_O_2_ and higher O_2_ concentration (Fig. 6A). Results were further verified by immunoblot analysis, which also showed a similar trend (Fig. 6B). The above results indicate that EhPKDL is stress responsive and may have some role to play in stress signaling. PKDs are often activated by upstream PKCs (1, 5, 16) by phosphorylation.

Therefore, immunoblot detected with anti-phospho-Ser/Thr/Tyr antibody showed EhPKDL is indeed phosphorylated on stress activation (Fig. 6B). On post-translational modification, analysis of the protein sequence identified a conserved ^137^SGK^139^, which is a PKC phosphorylation site. MEK and MAPK are proteins that are established stress signaling proteins, have also been checked for their mRNA expression. Transcriptional profile of EhMEK was found similar to EhPKDL, where its expression was found maximum during heat stress and was also seen to over-express for other stress conditions as well (Figure S4A).

There was no significant difference in expression in the case of EhMAPK (Figure S4B), which is well known to get activated by upstream MEKs by phosphorylation (18). Since stress activates EhPKDL, therefore, the protein was localized in different stress-activated cells. Images indicate that EhPKDL was highly expressed in stress-activated cells, and they are mostly seen towards the membrane periphery with a substantial amount in the cytoplasm as well (Fig. 6C)

Heat stress seems to activate EhPKDL many times more compared to any other stress. Therefore, further studies were carried out with heat stress alone. Since actin was identified as substrates of EhPKDL, heat shocked cells were co-localized for the EhPKDL and actin. EhPKDL, along with EhActin, was found to localize beneath the plasma membrane on heat stress, which otherwise also remain in the cytoplasm. On specific dsRNA treatment, there was reduced EhPKDL localization in membrane and cytoplasm. However, on specific dsRNA treatment, cells showed reduced colocalization of EhActin and EhPKDL (Fig. 7). Since actin are proven substrates of EhPKDL (Fig. 2B and 2C), now with their co-localization studies in heat stressed *E. histolytica*, reiterates EhPKDL’s involvement in actin rearrangement near the cell membrane.

**Figure. 6.**
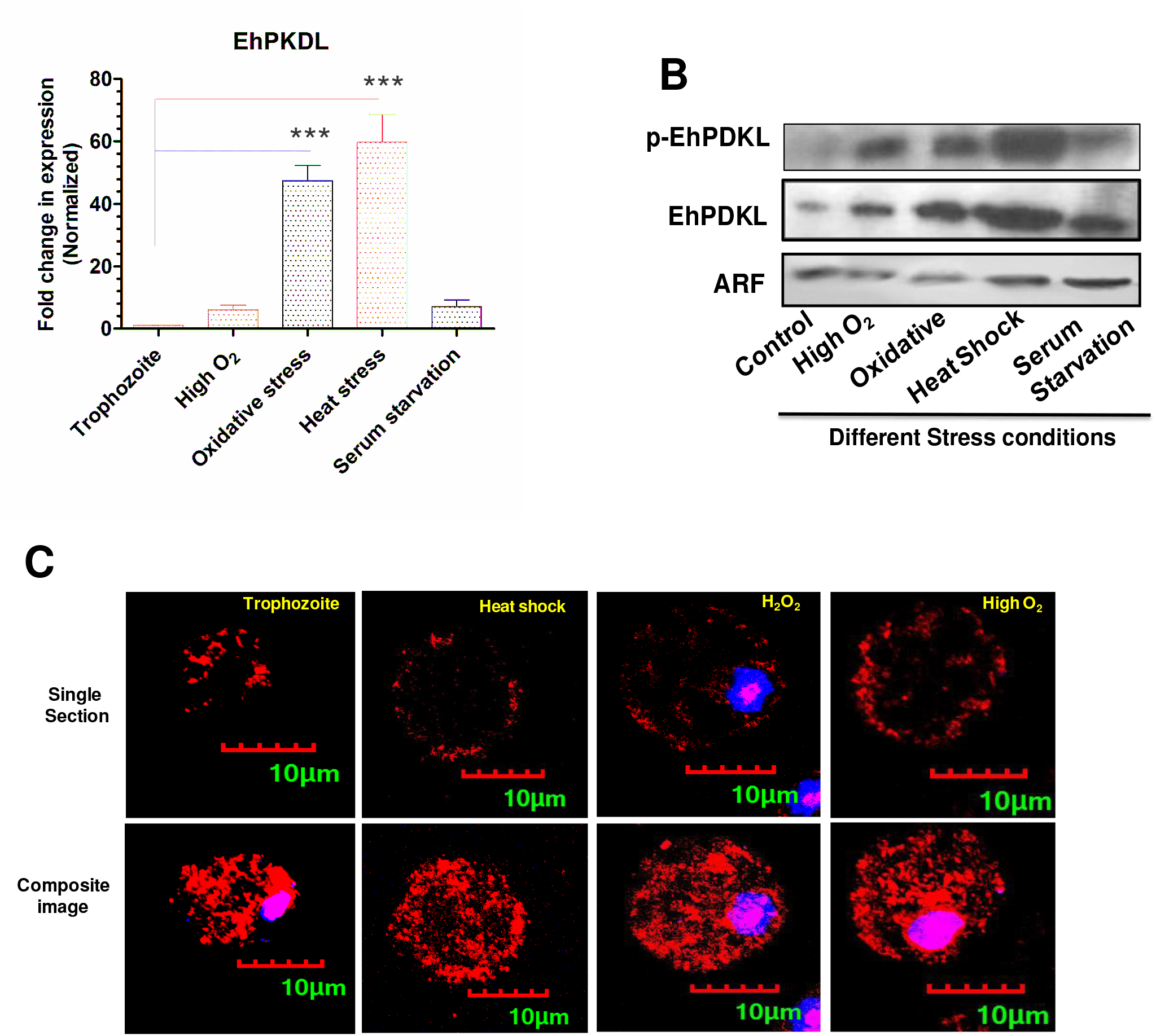
EhPKDL responds to stress and gets phosphorylated. A. *E. histolytica* was exposed to different stress conditions to estimate EhPKDL’s relative expression mRNA was analyzed using RT-PCR, B. Stimulated cells with different stress conditions shows higher levels of EhPKDL phosphorylation, as probed with anti-Ser/Thr/Tyr antibody followed by secondary anti-mouse IgG antibody. ADP-ribosylation factor (ARF) protein was considered as the internal control. C. Confocal micrograph stained for EhPKDL in stressed cells reaffirmed higher expression of EhPKDL, but now upon stress, EhPKDL was found to relocalize from cytoplasm to cell membrane in a significant amount. EhPKDL was stained with anti-EhPKDL antibody followed by TRITC labeled ant-rabbit antibody. DAPI stained the nucleus.

**Figure. 7.**
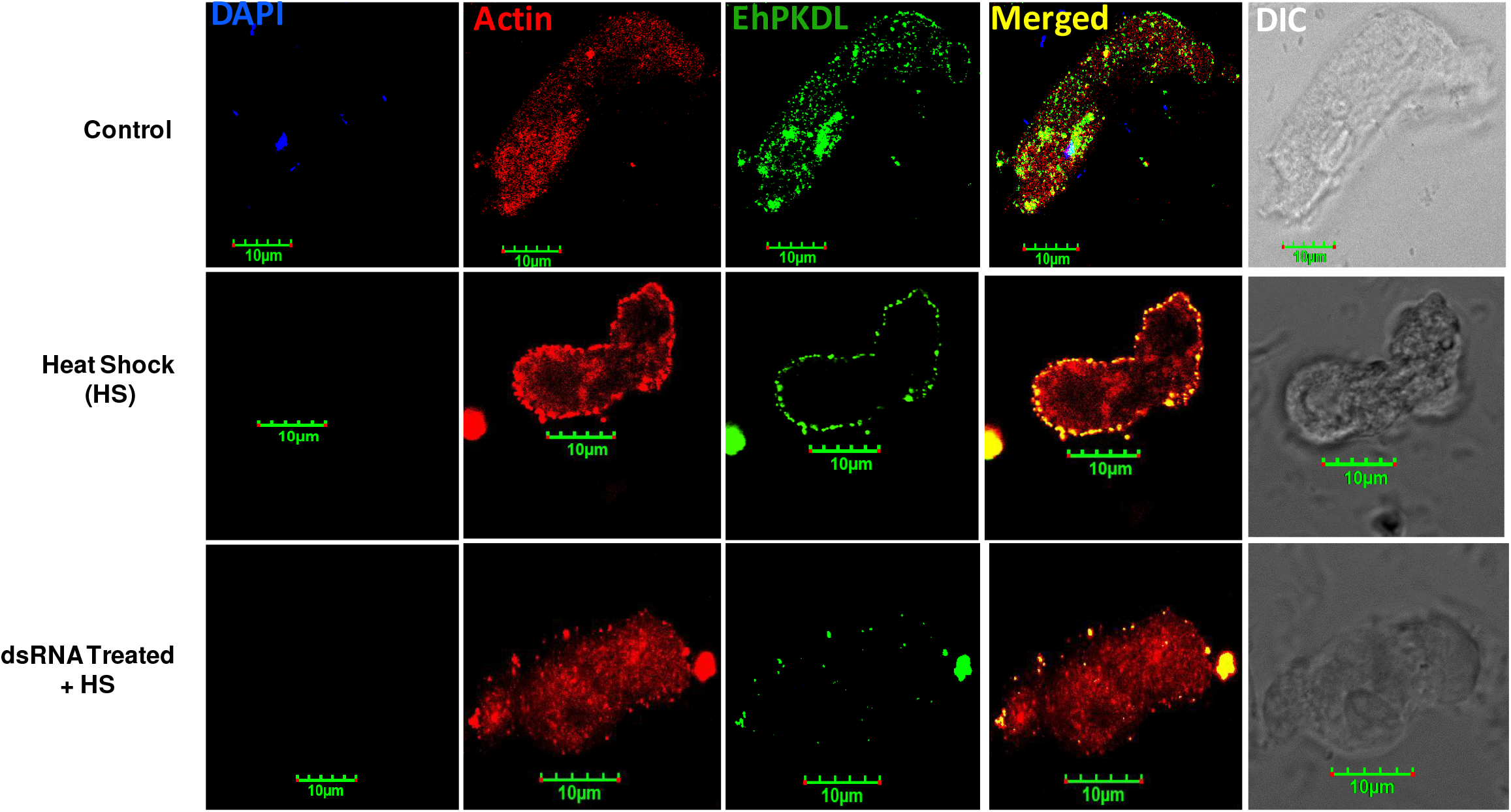
EhPKDL co-localizes with actin on heat stress. *E. histolytica* was stimulated on exposure to heat stress at 42°C for 1 h. Cells were then fixed and immunostained with anti-EhPKDL antibody followed by Alexa 488 labeled ant-rabbit antibody. Cells were stained for F-actin using Rhodamine-Phalloidin and DAPI stained nucleus. Finally, heat treated or untreated cells with or without soaking in EhPKDL dsRNA were analyzed under the confocal microscope. Microscopic analysis shows relocalization of EhPKDL from the cytoplasm to membrane along with increased F-actin aggregation beneath the cell membrane in heat stimulated cells, but not in EhPKDL specific dsRNA treated Eh cells.

## Discussion

Actin polymerization is sacrosanct to the cellular functioning, and a plethora of proteins are involved in manifesting its dynamism. Just in recent past an essential modulator of actin dynamics in *Eh viz*. Eh alpha kinase 1 (EhAK1), was reported to assists in actin polymerization (15). EhAK1 was also implicated in the recruitment of EhARPC1, a subunit of the actin branching complex Arp 2/3 at the phagocytic initiation site (15). The study gives us some molecular insight into this complex process but how de novo actin seeding takes place at the site of F-actin formation in phagocytic cup formation in *Eh*, remains an enigma. We here report a novel protein kinase as an actin nucleator which promotes an F-actin formation.

The protein identified was similar to Protein kinase D, with a kinase domain and Pleckstrin homology (PH) domain in common, but without cysteine-rich domains and so it is referred as PKD like kinase (EhPKDL). Since *Entamoeba* is an early branching eukaryote, regulatory domains could have evolved for stringent regulation and specific mode of activation in different tissue types in higher eukaryotes. Moreover, there is no similar PKD found in *Eh*, so this reported EhPKDL could supplement the role of PKDs in *Eh*. Also, the enzymatic analysis revealed that EhPKDL, like PKDs, is insensitive to staurosporine (19) and genistein. This novel PKD like kinase in *Eh* has unique substrate in the form of actin, which is the first report for this group of kinases. But what is known, is that PKD regulates the actin dynamics through other mediators like SSH1L, PAK4, cortactin, E-Cadherin, SNAIL, and RIN1 (20–27). EhPKDL bypasses regulators, and itself modulates actin polymerization in *Eh*, a departure from its conventional role. Binding of EhPKDL to monomeric actin is mediated by a G-actin binding site (^89^NVLFSADFEETSALGVYQSTVE^111^) similar to Depactin (10-27 residues) from starfish (28). On further analysis, EhPKDL was found to polymerize G-actin into a filamentous F-actin. Gel filtration profile indicates EhPKDL readily forms a dimer, and fluorescent quenching studies show molar stoichiometry of EhPKDL-actin interaction is 1:1. Therefore, we expect dimeric EhPKDL can at least absorb two G-actin molecules at a time. This pattern of monomeric actin interaction is well documented for actin nucleation proteins like Spir (29, 30) and Arp2/3 (31–34). We next tested whether the actin-binding sequences in EhPKDL participate in actin filament nucleation. The spatial arrangement of seed actin monomer in the nucleus is of extreme importance as it can determine the stability and length of F-actin (35). Since, EhPKDL predominantly exists in dimeric form, as inferred from gel filtration profile probably due to the PH domain. Thus, being a dimer, it has enriched its actin binding sites as a nucleating species. During nucleation, initially, it shows no difference in fluorescence intensity upon EhPKDL saturation with actin as compared to pyrenated G-actin alone. But on titration of unlabelled G-actin, the fluorescence intensity in the range of polymerized actin kept on increasing until the concentration crossed 1:1 ratio of EhPKDL: actin. This indicates that once actin binding site in EhPKDL is saturated, there can be a change in the configuration of actin-bound EhPKDL, which orients actin longitudinally. This arrangement may expose actin-actin interacting sides of bound actin, where newly arrived actin will come and fit stepwise to elongate into a stable F-actin. Confocal micrograph of early phagocytic cup showed the minuscule amount of colocalization between EhActin and EhPKDL, but not in the matured phagosome, where F-actin forms a wreath around the structure. This study also indicates that initial phagocytic cup requires *de novo* actin nucleation, but once stable F-actin formation is completed, EhPKDL withdraws from the structure. Until actin bound EhPKDL structure is solved, the hypothesized mode of actin nucleation by EhPKDL can be an attractive model but other assisting actin-binding protein in the nucleation of actin polymerization in *E. histolytica* cannot be ruled out. EhPKDL other than nucleation also phosphorylates G-actin *in vitro*. In case of *E. histolytica*, it has been reported that G-actin phosphorylated at Thr107 forms a stable F-actin (15), but for other organism phosphorylation of monomeric G-actin leads to inhibition of F-actin formation, including an amoeba *viz*. *Amoeba proteus* (36) and *D. discoideum*. In *D. discoideum*, Tyr-53 phosphorylation inhibits nucleation and elongation (37). Nucleation is a rate-limiting step in actin polymerization, and therefore, EhPKDL regulates all the cellular activities that are dependent on actin polymerization. Phagocytosis, capping, cell motility are few of the functions mediated by actin dynamics in *E. histolytica* that were studied in EhPKDL downregulated cells to assess in vivo importance of EhPKDL. All these processes were severely affected due to the lack of sufficient EhPKDL in dsRNA treated *E. histolytica* cells.

Erythrophagocytosis study in EhPKDL downregulated cells shows a reduced intake of RBCs. From the previous study, we know that attachment of RBC to the membrane recruits EhC2PK at the attachment site (Ca^2+^ dependent). EhCaBP1 then arrives at the site and interacts with EhC2PK (Ca^2+^ independent). By this time, some actin molecule begins to appear at the site along with EhCaBP3. Now, EhAK1 is brought to cupping site by binding with EhCaBP1 (Ca^2+^ dependent) (15), which also recruits Arp 2/3 complex for actin branching (15). We presume, G-actin at this site is nucleated with the help of EhPKDL, and maybe together with EhAK1, EhPKDL rapidly polymerizes the actin that helps form the cup. The rest of the process with the recruitment of Ehmyosin 1B through EhCaBP3 which stabilizes the cup may follow. But if the EhPKDL is in conduit with the above cascade of events, involved in phagocytosis, or it follows an entirely different pathway with another set of participating proteins is remain to be seen.

As far as capping is concerned EhPKDL downregulated *E. histolytica* cells could not successfully form a cap, and thus the ConA remain spread over the cell membrane. This clustering of surface glycoproteins followed by renewed refilling from the internal pool is anticipated due to actin dynamics. So when actin nucleation is aberrant due to deficiency of EhPKDL, cells were left with reduced cap formation. Similarly, cell motility is also mediated by actin polymerization, which helps in polarization, quintessential for amoebic movement.

Polarized *E. histolytica* usually has pseudopodia and a terminal uroid region. Pseudopodia give the direction to the cell along which come the cell mass with an intracellular churning of protoplasm called cell cyclosis regulated by self-generated chemokines and chemorepellents (38). On EhPKDL specific dsRNA treatment, cells lose its motility significantly due to reduced F-actin for the cell to polarize. EhPKDL is found to be stress responsive, and it becomes activated *via* phosphorylation along with its increased expression. Since EhPKDL was not found to autophosphorylate itself in *in vitro* kinase assay, therefore, we presumed it might be due to upstream activation by other kinases. Usually, PKD was found to be activated by phosphorylation by upstream PKCs, and Eh does have putative PKCs, so the activation of EhPKDL by PKCs cannot be ruled out. Moreover, *in silico* post-translational modification analysis of the sequence revealed a conserved ^137^SGK^139^ site where usually upstream PKCs phosphorylates PKDs and activates them. Thus, EhPKDL reported here is critical to cell functioning as it regulates actin modulation, thereby influencing all cellular functions mediated by actin dynamics (Fig. 8).

**Figure. 8.**
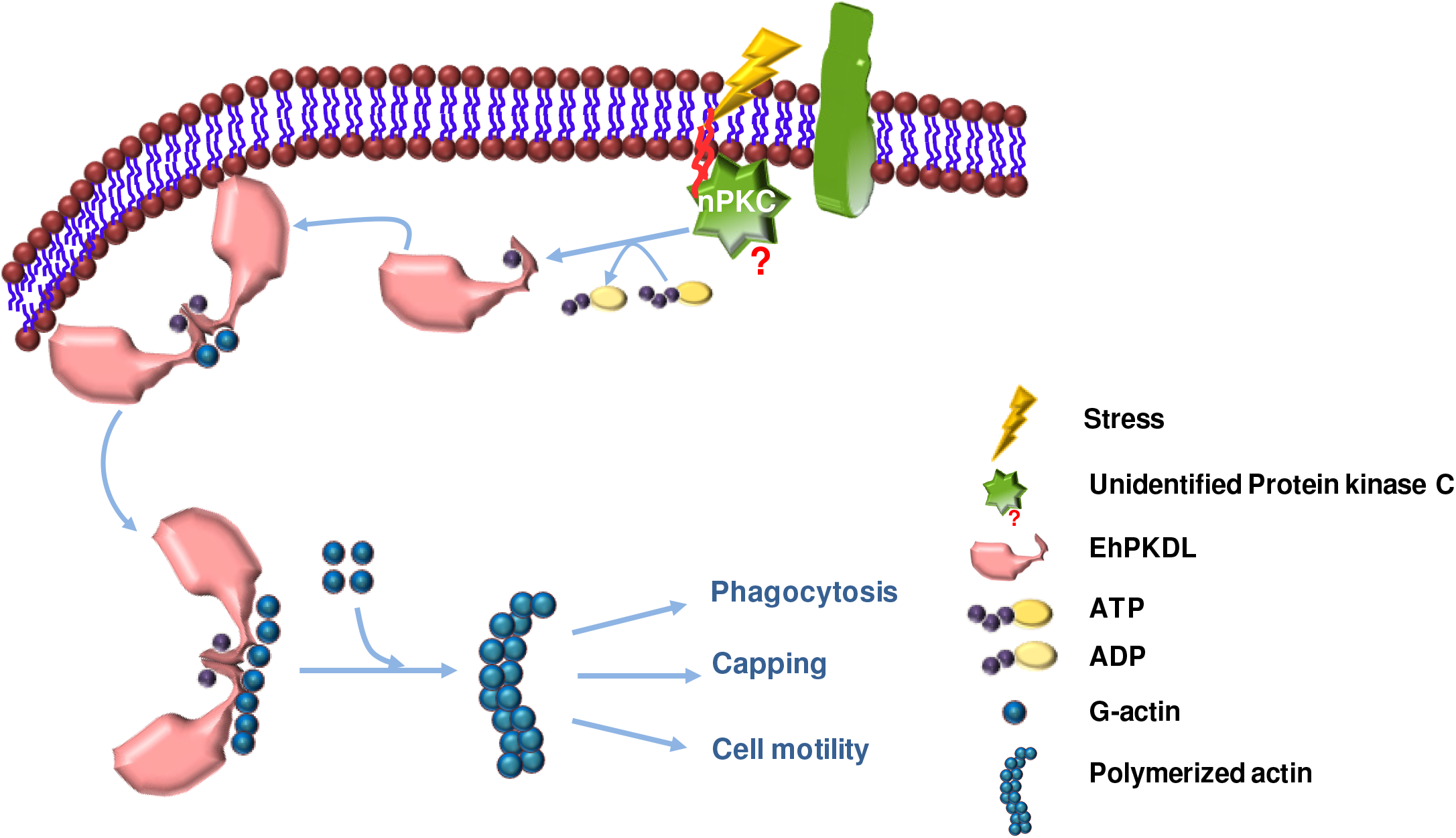
Schematic representation of EhPKDL’s mode of action in *E. histolytica*. In response to different kinds of stress, EhPKDL is activated upon on phosphorylation by a presumptive upstream Protein kinase C. Activated EhPKDL forms dimer and now is able to nucleate G-actin and hastens the process of actin polymerization. Thus, EhPKDL can effectively control various cellular processes dependent on action dynamics.

## Experimental procedures

### Cloning, expression and purification of EhPKDL

Full-length *Eh* antigenic protein was cloned into the pGEX-4T-1 vector with a GST-tag. The primer used had *Bam*HI site in forward primer and *Sal*I site in reverse primer. Vector was also digested with the same enzymes to generate compatible sticky ends. Positive clones were confirmed by fragment release on double digestion with the same combination of restriction enzymes. The protein now was expressed in BL21(DE3) *E. coli* strain after inducing with 0.2 mM IPTG at 16°C overnight. Proteins were initially affinity purified with immobilized glutathione-agarose beads. Bacterial cells lysed with buffer containing 50 mM Tris-Cl (pH 7.5), 150 mM NaCl, 5 mM EDTA, 1 mM 2-mercaptoethanol, 0.15 mM PMSF and 0.1% Triton X-100. Sonicated lysate was then allowed to bind to the pre-equilibrated glutathione-agarose beads, with the above buffer. Non-specifically bound proteins were removed by the wash buffer having 1XPBS (pH 7.4), 5 mM EDTA, 0.15 mM PMSF. Finally, GST-tag protein was eluted with the elution buffer having 100 mM Tris-Cl (pH 8.0) and 10 mM glutathione. Further proteins were purified to homogeneity in gel filtration chromatography using S-200 sepharose column (Amersham, USA). The composition of the buffered used in gel filtration was 10 mM Tris, 50 mM NaCl, and 2 mM DTT. The purity of EhPKDL (40 kDa) was assessed by separating the eluted fraction in SDS-PAGE.

### Enzyme kinetics

Phosphorylation of the substrate was measured by the incorporation of radioactive γ-^32P^ from [γ-^32P^]ATP. The 20 µl of phosphorylation reactions were performed at 30°C for 1 h in 50 mM HEPES buffer, pH 7.5, containing 50 mM NaCl, 1 mM DTT, 0.02% BSA, 10 mM MgCl2, 0.2 mM sodium vanadate and 3 µg of histone type III protein. For measuring Michaelis-Menten parameters varying [γ-^32P^]ATP concentrations were used (1, 2, 3, 5, 10, 25, 50, and 100 µM). For actin as substrate concentration was varied from 1, 2, 3, 5, 10, 25 and 50 µM and [γ-^32P^]ATP was at a final concentration of 2 µM. The assays were initiated by the addition of 10 nM recombinant kinase. For all kinase assays, 20 µl of 75 mM phosphoric acid was mixed with the reaction to stop the reaction and then applied to P81 phosphocellulose paper (2X2 cm). A minute after the incubation, the papers were washed thrice with 1 liter of fresh 75 mM phosphoric acid for 10min, then rinsed with 50 ml of acetone, and placed in the hood (5 min) to dry. The specific radioactivity of 32P-labeled substrates (cpm pmol^-1^) was determined by a scintillation counter from the amount of radioactivity detected from the known amount of total substrate that was applied to the P81 paper. The µM amount of phosphorylated product formed was determined by reference to the specific radioactivity of [γ-^32P^]ATP and the volume of the aliquot removed for quenching (20 µl). Initial velocities (v, µM s^-1^) were measured under conditions where total product formation represented 10% of the initial concentration of the limiting substrate. Negative control assays were carried out in conditions, where either the enzyme, substrates or [γ-^32P^]ATP were removed and in one case 2 mM EDTA was added. Staurosporine (20 µM), a Ser/Thr protein kinase inhibitor and genistein (50 µM), specific tyrosine kinase inhibitor were used to identify the enzyme type.

### Co-immunoprecipitation of interacting proteins

Cell lysis buffer for immunoprecipitation contained 10 mM Tris-HCl, pH 7.4, 150 mM NaCl, 2 mM β-ME, 50 mM NaF, 1 mM sodium orthovanadate, 2.5 mM sodium pyrophosphate, 1 mM β-glycerophosphate, 1 mM phenylmethylsulfonyl fluoride (PMSF), protease inhibitor cocktail, 1% Triton X-100 and 25 µM adenylyl-imidodiphosphate (AMP-PNP). After clear cell lysate was obtained, cellular debris was removed by centrifugation at 15,000 rpm for 15 min. The conjugated Protein A-Sepharose beads were incubated with *E. histolytica* lysate for 3 h at 4°C for pre-clearance. The pre-cleared lysate was then incubated with 30 µl of purified EhPKDL specific antibody, developed in rabbit, for overnight at 4°C. Fresh conjugated Protein A-sepharose was added and was incubated for 3 h in 4°C. The beads were then washed with wash buffer (10 mM Tris-Cl pH 7.4, 150 mM NaCl, 2 mM β-ME, 0.1% Triton 100 and protease inhibitor cocktail) thrice. The washed pellet was suspended in SDS polyacrylamide gel electrophoresis (PAGE) buffer and was boiled for 5 min and finally resolved by SDS-PAGE. The gel was stained by Coomassie brilliant blue R-250 dye following slandered protocol.

### Mass spectrometry of tryptic digested peptide

Specific silver-stained protein bands were cut out for in-gel trypsin digestion. Excised bands were further cut into tiny pieces. Distaining was done by repeated washing with 50 mM and 50% acetonitrile and finally with 100% acetonitrile. The reduction was made using 10 mM dithiothreitol solution, prepared in 100 mM NH_4_HCO_3_, at 56°C for 30 min. The above solution was removed, and 55 mM iodoacetamide (IAA) solution prepared in 100 mM NH_4_HCO_3_ was added and incubated further in the dark at room temperature, for 30 min. Tryptic digestion was done overnight at 37°C in the reaction mixture containing trypsin (10 ng µl^-1^ of 25 mM NH_4_HCO_3_). The digested peptides were mixed with a saturated solution of α-Cyano-4-hydroxycinnamic acid in 50% acetonitrile/H_2_O with 0.1% trifluoroacetic acid and 1.5 µl solutions were deposited on the MTP 384 ground steel plate. Finally, the sample was analyzed using MALDI-TOF MS (peptide mass fingerprinting). MALDI-TOF-MS analysis was carried out with Ultraflex TOF/TOF (Bruker Daltonics, Bremen, Germany) mass spectrometer, in positive ion reflectron mode. The spectral data were processed by Bruker Daltonics FLEX analysis software version 3.0. For external calibration, a standard peptide mixture (P.N: 206195, Bruker peptide calibration standard) was used.

### Purification and storage of *Entamoeba* actin

*Entamoeba* actin (EIN_089410) cloned in pET28a(+) vector expressing in Rosetta(DE3)pLysS was already available. Actin was initially purified from the IPTG (0.5mM) induced bacteria, by metal affinity (Ni-NTA) chromatography. Bacterial cells were lysed by sonication in a buffer containing 2 mM Tris-Cl (pH 8.0), 150 mM NaCl, 0.15 mM PMSF (Buffer 1). Sonicated lysate was then allowed to bind to the pre-equilibrated Ni-NTA agarose beads, with the Buffer 1. Non-specifically bound proteins were removed by the same buffer having additional imidazole in lower concentrations (20 mM and 100 mM) and finally, the 6xHis tagged actin was eluted by Buffer 1 having 300 mM imidazole. The affinity purified actin was dialyzed for 24 h in a buffer containing 2 mM Tris (pH 8.0), 0.1 mM CaCl_2_, 0.5 mM DTT so that all the imidazole is removed. Further, the actin was purified by gel filtration on Sephadex S-200 column in a buffer containing 2 mM Tris (pH 8.0), 0.1 mM CaCl_2_, 0.5 mM DTT, 0.2 mM ATP and 1 mM Sodium Azide. Purified actin was stored in the gel filtration buffer devoid of ATP and DTT.

### Pull-down assay

Proteins used for pull-down assays were previously purified in 20 mM Na-HEPES (pH 7.5), 200 mM NaCl, 2 mM MgCl_2_, 0.01% NP-40 and 5% glycerol. 10 μg of EiActin was immobilized on Ni-NTA beads as bait. GST-EhPKDL was mixed with pre-immobilized bait proteins in Ni-NTA beads in different concentrations (10, 20, 30, 50, 100 and 200 μg). Post incubation at 4°C on a turning wheel, extensive washing was done. Following which proteins were eluted by heating at 100°C for 5 min in a buffer containing 200 mM Tris-HCl (pH 6.8), 8% SDS (w/v), 40% glycerol, 0.4% bromophenol blue, and 100 mM DTT, and analyzed by SDS-PAGE with Coomassie R-250 staining. A negative control was maintained with only GST tag in both the experiments.

### Actin polymerization assay

Polymerization of actin was analyzed by fluorescence spectroscopy as per manufacturer’s protocol (cytoskeleton, USA). The assays were carried out in a Perkin Elmer fluorometer (LS55). A typical 100 μl polymerization reaction containing 3 μM G-actin (10% pyrene labeled G-actin), was saturated with EhPKDL or only GST protein at 2.5 μM and the reactions were carried out in polymerization buffer containing 5 mM Tris-HCl, pH 7.5, 1 mM dithiothreitol, 0.1 mM CaCl_2_, 0.01% NaN_3_, 100 mM KCl and 2 mM MgCl_2_, 1 mM ATP. Increase in fluorescence of pyrene-labeled actin (Cytoskeleton, USA) was measured with excitation at 366 nm and emission at 407 nm.

For actin nucleation studies unlabeled actin was isolated following routine purification protocol, and pyrenylated actin was sourced from Cytoskeleton, USA. Aliquots of 0.2 µM of pyrene actin was introduced stepwise to a solution containing EhPKDL (0.2 µM) in 800 μL buffer having buffer (10 mM imidazole, pH 7.2, 2 mM MgCl_2_, 0.2 mM CaCl_2_ and 1 mM ATP). The mixture was then incubated for 3 min at room temperature before starting the scan. Unlabelled actin was subsequently titrated in 0.4 μM aliquots until desirable polymerization was obtained. Fluorescence emission was analyzed between 350 and 500 nm at an excitation of 342 nm (28).

### Preparation of dsRNA construct for EhPKDL silencing

The region of the gene used for dsRNA production was chosen as such that it was least similar to other *E. histolytica* but contained the highest possible of number of siRNAs (39). The 189 bp region from 122 bp to 311 bp of the EhPKDL gene was found most suitable. A non-specific dsRNA, a 150 bp region of oxygen-independent flavin mononucleotide (FMN)-based fluorescent protein of *Bacilus subtilis* (BS1) was chosen as a negative control. These gene fragments were PCR amplified with gene-specific primers (Table S1) and sub-cloned into the TA-cloning vector pTZ57R/T (Fermentas, USA). DNA inserts were cut out from the TA-cloning constructs using restriction enzymes (*Bgl*II/*Xho*I for EhPKDL and BS1) and were subsequently cloned into (*Bam*HI/*Xho*I) double digested expression vector pL4440 (Addgene, USA) which is flanked by T7 promoters at both ends. The plasmid constructs made were verified by restriction digestion and DNA sequencing. The plasmids construct were transformed and expressed in HT115 (DE3) *E. coli* strain, which is RNase III-deficient. For bacterial transformation, competent cells were prepared in calcium chloride (41).

Competent cells transformed with recombinant plasmids were selected on LB plates containing ampicillin (100 µg ml^-1^) and tetracycline (12.5 µg ml^-1^). For dsRNA expression, overnight bacterial cultures of confirmed recombinant clones were inoculated in 500 ml LB broth supplemented with ampicillin and tetracycline at 37°C with shaking at 220 rpm. Bacterial cells were induced for dsRNA production with 0.5 mM IPTG when the optical density of the culture reached 0.4 at 600 nm.

### Purification of dsRNA

Expressed dsRNA in HT115(DE3) *E. coli* strain were isolated using a previously described method (41, 42). The bacterial pellet was suspended in 1/20th of the initial induction volume containing 10 mM EDTA, 1 M ammonium acetate. An equal volume of phenol: chloroform (1:1) mixture, was added and incubated at 60°C for 30 min, followed by centrifugation at 9300Xg for 15 min. The upper aqueous layer was separated and mixed with an equal volume of isopropanol and incubated at −20°C for 30 min. The nucleic acid pellet was obtained after centrifugation at 9300g for 30 min. The pellet was further washed with 70% ethanol, air-dried, and finally resuspended in 5 ml of nuclease-free water. For removal of contaminating DNA and single-stranded RNA, the nucleic acid mixture was treated with 0.1 U l^-1^ of DNase (Fermentas) and 0.2 g l^-1^ of RNase A (Sigma) at 37°C for 3 h.

Further purification of dsRNA was obtained with Ribozol (Ameresco, USA) chloroform treatment (as suggested by manufacture’s protocol) and kept at room temperature for 2–3 min. Aqueous layer separated after centrifugation at 16,000 × g for 15 min was collected and mixed with an equal volume of isopropanol and incubated at −20°C for 30 min. Purified dsRNA pellet was obtained on centrifugation at 10,000 rpm for 25 min at 4°C. The pellet was then washed twice with 70% ethanol at 9300 × g for 5 min at 4°C. The pellet was then air dried and dissolved in 1 ml of nuclease-free water. Isolated dsRNAs were analyzed by agarose gel electrophoresis, and their concentrations were determined using a spectrophotometer (Thermo Scientific, USA).

### Silencing of EhPKDL with specific dsRNA

Exponentially growing *E. histolytica* cells were harvested and seeded in fresh TYI-S-33 media at a final density of 5 × 10^5^ cells ml^-1^ were soaked in purified 200 µg ml^-1^ of EhPKDL specific dsRNA for 24 h at 25°C. *E. histolytica* cells were then harvested and analyzed for EhPKDL silencing by sqRT-PCR and western blot.

### sqRT-PCR analysis

For sqRT-PCR analysis for studying gene expression, total RNA was isolated from *E. histolytica* cells using Ribozol (Ameresco, USA) following the manufacture’s protocol. The total RNA isolated was treated with RNase-free DNase (Fermentas, USA) to remove the contaminating DNA. Approximately 1 µg of total RNA was used for cDNA synthesis using the MMLV reverse transcriptase (Fermentas, USA) following the manufacturer’s protocol. For sqRT-PCR, gene-specific primers were designed to amplify a small region (200–400 bp) from the 3’ end of the genes (Appendix). Usually, a PCR mixture of 50 µl was prepared. The reaction cycle typically used was 94°C for 1 min, followed by 22–30 cycles (depending on the saturation of the amplified products) at 94°C for 30 s, 50°C-55°C for 30 s, and 72°C for 30 s, followed by a final extension at 72°C for 10 min. Positive and negative controls with genomic DNA and total RNA, respectively, without RT, were used as templates. ADP-ribosylation factor (ARF) was used as an internal control to normalize the initial variations in sample concentration and variations in reaction efficiency from sample to sample. PCR products were analyzed on 2% agarose gels with ethidium bromide (10 g ml^-1^) and photographed with UV light box. For quantification, the pixel intensity of the individual band was determined using ImageJ (NIH Image, Bethesda, MD). The experiments were repeated at least thrice, and the means (±standard deviations) of three replicates were presented as a relative mRNA expression value. The sqRT-PCR was standardized on parameters, such as primer T_m_ and concentration, MgCl_2_ concentration, and the number of cycles for each gene was optimized for obtaining most accurate results (43).

### Measuring gene expression by RT-PCR

Total RNA from cells under different stress conditions was prepared with TRIzol Reagent (Thermo Fisher Scientific) as recommended by the manufacturer and treated with DNase I (Fermentas, USA) until free of DNA. Absence of DNA in the samples was confirmed by 40 cycles of PCR with actin primers with no detectable PCR band on agarose gel electrophoresis. 3µg of mRNA was then reverse transcribed using Superscript II using oligodT primers (Invitrogen). Reverse transcriptase (RT)-PCR was performed at 42°C for 50 min. Real-time PCR was carried out in 0.2 ml microfuge tubes in a reaction volume of 10 µl using the device Eppendorf MasterCycler RealPlex (Thermo Fisher Scientific) and PowerUp™ SYBR® Green Master Mix(Thermo Fisher Scientific) with 0.3 µM primers and 200ng of cDNA. The default PCR Cycling conditions were employed: 2 min at 50°C for UDG activation, 10 min at 95°C for AmpliTaq DNA Polymerase activation followed by 40 cycles of 15 s at 95°C, 1 min at 60°C. Melting curve analysis was done to confirm the specificity of amplicons. Quantification of gene expression differences was performed using the ‘delta-delta-CT’ method with ARF as the normalizer gene. All experiments were carried out at least in triplicate with templates from independent experiments.

### Immunobloting

For expression analysis with western blot, 60 µg of total cell lysate (lysate was prepared as discussed in section 3.2.3 in materials and methods) estimated by Bradford assay kit (Genei, India) using BSA as a standard, were separated on 10–12% SDS–PAGE. The protein was then transferred on to a polyvinylidene fluoride membrane (PVDF) (Millipore, USA) using a semi-dry transfer system (Bio-Rad, USA). The membranes were then blocked for non-specific antibody binding with 5% skimmed milk (HiMedia, India) and for specific detection of phosphor-proteins 3% BSA (HiMedia, India) in 1X phosphate-buffered saline with 0.1% Tween-20 (PBST), pH 7.4 was used. The individual proteins were detected with specific polyclonal antibodies raised in rabbit and antibody dilution used were anti-EhPKDL, 1:3000; anti-ARF, 1:2000; Ser/Thr/Tyr mouse monoclonal antibody (Abcam, USA), 2 µg/ml. Binding with primary antibody carried out in room temperature for 1 h. After intermittent washing with PBST binding with secondary anti-rabbit immunoglobulins antibody conjugated to HRP (1:5,000, Sigma, USA) was allowed for about 1 h at room temperature. Finally, blots were developed by ECL reagents (Millipore, USA).

### Phagocytosis of red blood cells by *E. histolytica*

Human red blood cells (RBC) were collected from blood, were washed with 1XPBS followed by incomplete TYI-S-33 for three times each. Log phase growing *E. histolytica* cells treated or untreated with EhPKDL dsRNA for 24 h, were harvested in 1XPBS. Harvested *E. histolytica* cells were incubated with RBCs in 1:10 ratio at 37°C for 5 and 10 mins. Total cells were collected by centrifugation, and non-phagocytosed RBCs were lysed with cold distilled water. *E. histolytica* cell with engulfed RBCs was separated by centrifugation at 1000 g for 2 min and subsequently washed twice. The cell pellet was finally resuspended in 1 ml formic acid to burst *E. histolytica* cells containing engulfed RBCs. The optical density of the samples was measured at 400 nm using formic acid as the blank (15). For confocal studies, *E. histolytica* cell lysis was avoided; rather cells were fixed in 4% paraformaldehyde (PFA).

### Induction of cap in *E. histolytica*

Capping in *E. histolytica* was induced as described by published methods (44). In short, log phase *E. histolytica* were harvested, washed with 1XPBS. Washed cells were treated with TRITC tagged Concanavalin A (20 µg ml^-1^) (Sigma, USA) for 1 h at 4°C. For cap induction cells were incubated at 37°C for 5 min. After Concanavalin A-TRITC treatment, *E. histolytica* cells were fixed with 3.7% PFA.

### Determination of *E. histolytica* motility

Motility of the *Eh* trophozoites was determined by using polycarbonate transwell membrane inserts (8-μm pore size, 6.5-mm diameter, HiMedia, India), as described by Shahi et al., (2016). Briefly, Eh cells untreated and treated with dsRNA (for 24 h) were harvested and then washed three times in serum-free Diamond’s TYI-S-33 media. In the meanwhile, 24-well cell culture plates were filled with serum-free Diamond’s TYI-S-33 media, and in each well, a transwell membrane was inserted. Now the 500-μl harvested trophozoites having 25×10^5^ trophozoites/ml were then loaded into the transwell membranes, and the plates were then placed in an anaerobic chamber for three hours at 37°C. After the incubation, the transwell membrane inserts were removed from the wells, and the migrated trophozoites attached to the bottom of the wells were counted for determining the extent of cell motility.

### Induction of stress in *E. histolytica*

The *E. histolytica* trophozoites grown for 48 h were subjected to different stresses. Cells were kept under heat shock at 42°C for 1 h; exposure to H_2_O_2_ was done at a concentration of 0.7 mM for about 1 h. Since *E. histolytica* is microaerophilic, higher oxygen does induce stress. Therefore, high O_2_ stress was induced by seeding *E. histolytica* cells in 25 cm^2^ culture flask having 5 ml TYI-S-33 media for 3 h. In another set of experiment, *E. histolytica* cells were cultured in serum-free TYI-S-33 media for 12 h at 37°C. Normal trophozoites grown for 48 h were taken as control.

### Immunofluorescence staining for microscopy

For RBC engulfed and all kinds of stress-induced *E. histolytica* cells, fixing were done in 3.7% pre-warmed PFA for 30 min. Fixed cells were permeabilized with 0.1% Triton X-100 in PBS for 1 min, and immediately cells were washed with PBS thrice and subsequently treated with 50 mM NH_4_Cl. Non-specific staining was avoided with pre-incubation in 1% BSA in PBS for 30 min. For immunostaining, cells were incubated with primary antibody for 1 h at 37°C followed by washing (thrice) with 1% BSA in PBS. After that, cells were treated with secondary antibody was 30 min at 37°C. The polyclonal anti-EhPKDL antibody used was raised in rabbit and subsequently purified with Protein A-Sepharose (Invitrogen, USA). Dilutions of antibody used were: anti-EhPKDL 1:500, anti-rabbit Alexa 488 at 1:300 dilution, anti-rabbit Alexa 405 at 1:200 dilution (Molecular Probes) and anti-rabbit TRITC at a dilution of 1:250 (Sigma). For staining actin, Rhodamine-Phalloidin and Alexa 488-Phalloidin (Molecular Probes) was used at a dilution of 1:250. FLUOVIEW FV1000 laser scanning microscope (Olympus, Japan) was used confocal imaging. Colocalization analysis was done by using online Olympus FLUOVIEW software. Post microscopy the images were processed further using FV10-ASW 1.7 viewer.

### Statistical analysis

Estimated data are expressed as the mean ± standard error. Statistical significance was ascertained with the unpaired two-tailed Student’s t-test and one-way ANOVA test. Significance in mean differences were designated as *pvalue≤0.05, **p-value≤0.01, ***p-value≤0.005. All statistical calculations were either done in Origin 8 (OriginLab), Microsoft Excel or GrpahPad Prism.

## Acknowledgement

We thank DST-FIST for confocal microscopy facility at Department Biotechnology, IIT Kharagpur. MB and SN are thankful to UGC and CSIR, Govt. of India for fellowship. SKG is thankful to DBT and MHRD, Govt. of India, and IIT Kharagpur for funding of the research project.

## Author Contributions

Conceived and designed the experiments: MB, SKG, AKD. MB and SN performed the experiments. SKG and MB wrote the manuscript.

## Abbreviations

AC: acidic domain
AP: apolar region
C1: DAG-binding domain that contains C1a and C1b domains
P: proline-rich region
PH: Pleckstrin homology
OSBP: oxysterol-binding protein
PI4KIIIβ: phosphatidylinositol 4-Kinase III-β
CERT: ceramide trafficking protein
RUNX2: runt-related transcription factor 2
MEF2: myocyte enhancer factor-2
HDAC: class IIa histone deacetylases
SSH1L: slingshot protein phosphatase 1
RIN1: Ras and Rab interactor 1
PAK4: p21-activating kinase 4
MAPK: mitogen activated protein kinase
MEK: mitogen-activated protein kinase kinase.

**Abbreviations of the organisms in the phylogenetic tree**
Cf: Carpodacus formosanus
Cg: Crassostrea gigas
Cj: Callithrix jacchus
Cm: Chelonia mydas
Cn: Carpodacus nipalensis
Cq: Culex quinquefasciatus
Cv: Carpodacus vinaceus
Dr: Danio rerio
Ec: Equus caballus
Hs: Homo sapiens
Mm: Mus musculus
Mp: Mustela putorius furo
Ps: Pelodiscus sinensis
Pt: Pan troglodytes
Rn: Rattus norvegicus
Sm: Schmidtea mediterranea
Ss: Sus scrofa
Tg: Taeniopygia guttata.

## References

1. Rykx A, De Kimpe L, Mikhalap S, Vantus T, Seufferlein T, Vandenheede JR, Van Lint J. 2003. Protein kinase D: A family affair. FEBS Lett 546:81–86.

2. Roy A, Ye J, Deng F, Wang QJ. 2017. Protein kinase D signaling in cancer: A friend or foe? Biochim Biophys Acta Rev Cancer 1868:283–294.

3. Lint J Van, Rykx A, Vantus T, Vandenheede JR. 2002. Getting to know protein kinase D. Int J Biochem Cell Biol 34:577–581.

4. Fu Y, Rubin CS. 2011. Protein kinase D: coupling extracellular stimuli to the regulation of cell physiology. EMBO Rep 12:785–796.

5. Steinberg SF. 2012. Regulation of Protein Kinase D1 Activity. Mol Pharmacol 81:284–291.

6. Ellwanger K, Hausser A. 2013. Physiological functions of protein kinase D in vivo. IUBMB Life 65:98–107.

7. Nhek S, Ngo M, Yang X, Ng MM, Field SJ, Asara JM, Ridgway ND, Toker A. Mol Biol Cell. 2010. Regulation of Oxysterol-binding Protein Golgi Localization through Protein Kinase D–mediated Phosphorylation. Mol Biol Cell 21:2327–2337.

8. Fugmann T, Hausser A, Schöffler P, Schmid S, Pfizenmaier K, Olayioye MA. 2007. Regulation of secretory transport by protein kinase D-mediated phosphorylation of the ceramide transfer protein. J Cell Biol 178:15–22.

9. Jensen ED, Gopalakrishnan R, Westendorf JJ. 2009. Bone morphogenic protein 2 activates protein kinase D to regulate histone deacetylase 7 localization and repression of Runx2. J Biol Chem 284:2225–2233.

10. Xu X, Ha C-H, Wong C, Wang W, Hausser A, Pfizenmaier K, Olson EN, McKinsey TA, Jin Z-G. 2007. Angiotensin II Stimulates Protein Kinase D–Dependent Histone Deacetylase 5 Phosphorylation and Nuclear Export Leading to Vascular Smooth Muscle Cell Hypertrophy. Arterioscler Thromb Vasc Biol 27:2355–2362.

11. Olayioye MA, Barisic S, Hausser A. 2013. Multi-level control of actin dynamics by protein kinase D. Cell Signal 25:1739–1747.

12. Chang JK, Ni Y, Han L, Sinnett-Smith J, Jacamo R, Rey O, Young SH, Rozengurt E. 2017. Protein kinase D1 (PKD1) phosphorylation on Ser^203^ by type I p21-activated kinase (PAK) regulates PKD1 localization. J Biol Chem 292:9523–9539.

13. Bencsik N, Sziber Z, Liliom H, Tarnok K, Borbely S, Gulyas M, Ratkai a., Szucs a., Hazai-Novak D, Ellwanger K, Racz B, Pfizenmaier K, Hausser a., Schlett K. 2015. Protein kinase D promotes plasticity-induced F-actin stabilization in dendritic spines and regulates memory formation. J Cell Biol 210:771–783.

14. Babuta M, Mansuri MS, Bhattacharya S, Bhattacharya A. 2015. The *Entamoeba histolytica*, Arp2/3 Complex Is Recruited to Phagocytic Cups through an Atypical Kinase EhAK1. PLOS Pathog 11:e1005310.

15. Mansuri MS, Bhattacharya S, Bhattacharya A. 2014. A Novel Alpha Kinase EhAK1 Phosphorylates Actin and Regulates Phagocytosis in *Entamoeba histolytica*. PLoS Pathog 10:e1004411.

16. Chávez-Munguía B, Talamás-Rohana P, Castañón G, Salazar-Villatoro L, Hernández-Ramírez V, Martínez-Palomo A. 2012. Differences in cap formation between invasive *Entamoeba histolytica* and non-invasive Entamoeba dispar. Parasitol Res 111:215–221.

17. Wang QJ. 2006. PKD at the crossroads of DAG and PKC signaling. Trends Pharmacol Sci 27:317–323.

18. Ghosh AS, Ray D, Dutta S, Raha S. 2010. EHMAPK, the mitogen-activated protein kinase from *Entamoeba histolytica* is associated with cell survival. PLoS One 5.

19. Zhang W, Zheng S, Storz P, Min W. 2005. Protein kinase D specifically mediates apoptosis signal-regulating kinase 1-JNK signaling induced by H_2_O_2_ but not tumor necrosis factor. J Biol Chem 280:19036–19044.

20. Du C, Zhang C, Hassan S, Biswas MHU, Balaji KC. 2010. Protein kinase D1 suppresses epithelial-to-mesenchymal transition through phosphorylation of snail. Cancer Res 70:7810–7819.

21. Eiseler T, Döppler H, Yan IK, Kitatani K, Mizuno K, Storz P. 2009. Protein kinase D1 regulates cofilin-mediated F-actin reorganization and cell motility through slingshot. Nat Cell Biol 11:545–556.

22. Eiseler T, Hausser A, De Kimpe L, Van Lint J, Pfizenmaier K. 2010. Protein kinase D controls actin polymerization and cell motility through phosphorylation of cortactin. J Biol Chem 285:18672–18683.

23. Jaggi M, Rao PS, Smith DJ, Wheelock MJ, Johnson KR, Hemstreet GP, Balaji KC. 2005. E-Cadherin Phosphorylation by Protein Kinase D1 / Protein Kinase C µ is Associated with Altered Cellular Aggregation and Motility in Prostate Cancer is Associated with Altered Cellular Aggregation and Motility in Prostate Cancer 483–492.

24. Nagel AC, Schmid J, Auer JS, Preiss A, Maier D. 2010. Constitutively active Protein kinase D acts as negative regulator of the Slingshot-phosphatase in Drosophila. Hereditas 147:237–242.

25. Peterburs P, Heering J, Link G, Pfizenmaier K, Olayioye M a., Hausser A. 2009. Protein kinase D regulates cell migration by direct phosphorylation of the cofilin phosphatase slingshot 1 like. Cancer Res 69:5634–5638.

26. Spratley SJ, Bastea LI, Döppler H, Mizuno K, Storz P. 2011. Protein kinase D regulates cofilin activity through p21-activated kinase 4. J Biol Chem 286:34254–34261.

27. Barišić S, Nagel AC, Franz-Wachtel M, Macek B, Preiss A, Link G, Maier D, Hausser A. 2011. Phosphorylation of Ser 402 impedes phosphatase activity of slingshot 1. EMBO Rep 12:527–533.

28. Mabuchi I. 1983. An actin-depolymerizing protein (depactin) from starfish oocytes: Properties and interaction with actin. J Cell Biol 97:1612–1621.

29. Ducka AM, Joel P, Popowicz GM, Trybus KM, Schleicher M, Noegel A a, Huber R, Holak T a, Sitar T. 2010. Structures of actin-bound Wiskott-Aldrich syndrome protein homology 2 (WH2) domains of Spire and the implication for filament nucleation. Proc Natl Acad Sci U S A 107:11757–11762.

30. Quinlan ME, Heuser JE, Kerkhoff E, Mullins RD. 2005. Drosophila Spire is an actin nucleation factor. Nature 433:382–388.

31. Machesky LM, Mullins RD, Higgs HN, Kaiser D a, Blanchoin L, May RC, Hall ME, Pollard TD. 1999. Scar, a WASp-related protein, activates nucleation of actin filaments by the Arp2/3 complex. Proc Natl Acad Sci U S A 96:3739–3744.

32. Blanchoin L, Amann KJ, Higgs HN, Marchand J, Kaiser DA, Pollard TD. 2000. Direct observation of dendritic actin complex and WASP / Scar proteins 171:1007–1011.

33. Higgs HN, Pollard TD. 2000. Activation by Cdc42 and PIP2 of Wiskott-Aldrich Syndrome protein (WASp) stimulates actin nucleation by Arp2/3 complex. J Cell Biol 150:1311–1320.

34. Yarar D, Wayne T, Arie A, Welch MD. 1999. The Wiskott-Aldrich syndrome protein directs actin-based motility by stimulating actin nucleation with the Arp2/3 complex. Curr Biol 9:555–558.

35. Bosch M, Le KHD, Bugyi B, Correia JJ, Renault L, Carlier MF. 2007. Analysis of the Function of Spire in Actin Assembly and Its Synergy with Formin and Profilin. Mol Cell 28:555–568.

36. Sonobe S, Takahashi S, Hatano S, Kuroda K. 1986. Phosphorylation of Amoeba G-actin and its effect on actin polymerization. J Biol Chem 261:14837–14843.

37. Liu X, Shu S, Hong M-SS, Levine RL, Korn ED. 2006. Phosphorylation of actin Tyr-53 inhibits filament nucleation and elongation and destabilizes filaments. Proc Natl Acad Sci U S A 103:13694–13699.

38. Zaki M, Andrew N, Insall RH. 2006. *Entamoeba histolytica* cell movement: a central role for self-generated chemokines and chemorepellents. Proc Natl Acad Sci U S A 103:18751–18756.

39. Reynolds A, Leake D, Boese Q, Scaringe S, Marshall WS, Khvorova A. 2004. Rational siRNA design for RNA interference. Nat Biotechnol 22:326–330.

40. Solis CF, Santi-Rocca J, Perdomo D, Weber C, Guillén N. 2009. Use of bacterially expressed dsrna to downregulate *Entamoeba histolytica* gene expression. PLoS One 4:e8424

41. Timmons L, Court DL, Fire A. 2001. Ingestion of bacterially expressed dsRNAs can produce specific and potent genetic interference in Caenorhabditis elegans. Gene 263:103–112.

42. Samanta SK, Ghosh SK. 2012. The chitin biosynthesis pathway in *Entamoeba* and the role of glucosamine-6-P isomerase by RNA interference. Mol Biochem Parasitol 186:60–68.

43. Marone M, Mozzetti S, De Ritis D, Pierelli L, Scambia G. 2001. Semiquantitative RT-PCR analysis to assess the expression levels of multiple transcripts from the same sample. Biol Proced Online 3:19–25.

44. Espinosa-Cantellano M, Chávez B, Calderón J, Martínez-Palomo A. 1992. *Entamoeba histolytica*: electrophoretic analysis of isolated caps induced by several ligands. Arch Med Res. 23:81–85.

